# Comprehensive quantification of the modified proteome reveals oxidative heart damage in mitochondrial heteroplasmy

**DOI:** 10.1101/296392

**Authors:** Navratan Bagwan, Elena Bonzon-Kulichenko, Enrique Calvo, Ana Victoria Lechuga-Vieco, Spiros Michalakopoulos, Marco Trevisan-Herraz, Iakes Ezkurdia, José Manuel Rodríguez, Ricardo Magni, Ana Latorre-Pellicer, José Antonio Enríquez, Jesús Vázquez

**Affiliations:** Centro Nacional de Investigaciones Cardiovasculares Carlos III (CNIC), Madrid, 28049; Spain; CIBER Cardiovascular Diseases (CIBERCV), Madrid, Spain; CIBERES: C/ Melchor Fernández-Almagro 3, 28029, Madrid, Spain; CIBERFES: C/ Melchor Fernández-Almagro 3, 28029, Madrid, Spain

**Keywords:** post-translational, modifications, mass spectrometry, proteomics, bioinformatics, heteroplasmy, oxidative phosphorylation, mitochondria

## Abstract

Post-translational modifications hugely increase the functional diversity of proteomes. Recent algorithms based on ultratolerant database searching are forging a path to unbiased analysis of peptide modifications by shotgun mass spectrometry. However, these approaches identify only half of the modified forms potentially detectable and do not map the modified residue. Moreover, tools for the quantitative analysis of peptide modifications are currently lacking. Here, we present a suite of algorithms that allow comprehensive identification of detectable modifications, pinpoint the modified residues, and enable their quantitative analysis through an integrated statistical model. These developments were used to characterize the impact of mitochondrial heteroplasmy on the proteome and on the modified peptidome in several tissues from 12-week old mice. Our results reveal that heteroplasmy mainly affects cardiac tissue, inducing oxidative damage to proteins of the oxidative phosphorylation system, and provide a molecular mechanism that explains the structural and functional alterations produced in heart mitochondria.

**Highlights:**

- Identifies all protein modifications detectable by mass spectrometry
- Locates the modified site with 85% accuracy
- Integrates quantitative analysis of the proteome and the modified peptidome
- Reveals that mtDNA heteroplasmy causes oxidative damage in heart OXPHOS proteins

## INTRODUCTION

Shotgun mass spectrometry (MS)-based proteomics (Link et al., 1999) has become a powerful tool for biotechnological and biomedical research. Advances in speed and sensitivity allow the generation of millions of spectra per experiment, but only a minority of these spectra can be mapped to proteins (Griss et al., 2016; Skinner and Kelleher, 2015). A large proportion of unassigned spectra are thought to arise from peptides containing sequence variants or unknown chemical and posttranslational modifications (PTM) (Griss et al., 2016), and their characterization is one of the most interesting and challenging goals of current proteomics. A number of computational methods have been proposed for the detection of these unmatched peptides (Berna et al., 2007; Griss et al., 2016; Kim and Pevzner, 2014; Ma and Lam, 2014; Shortreed et al., 2015; Zhang et al., 2009). Recently, an “open search” (OS) strategy, where precursor mass tolerances of hundreds of Da were used with a conventional search engine, was reported to identify modified peptides at an unprecedented scale (Skinner and Kelleher, 2015). Another report demonstrated the ability to perform OS at orders-of-magnitude faster speeds using a novel fragmentation-ion indexing algorithm (MSFragger) (Kong et al., 2017). Although these two methods may have a considerable impact on the field, opening the way to true hypothesis-free analysis of PTMs by MS, OS algorithms still rely on the chance that the modification leaves enough unaffected fragment ions for matching by the search engine (Figure 1A). Therefore OS strategies can identify only approximately half of the modified peptides detectable by conventional “closed” searches (CS) (Chick et al., 2015). Moreover, the modification site cannot be directly identified by existing OS approaches. In addition, OS methods have not previously been used to quantify PTM alterations, and a general statistical model for the analysis of data of this kind is currently lacking.

**Figure 1.**
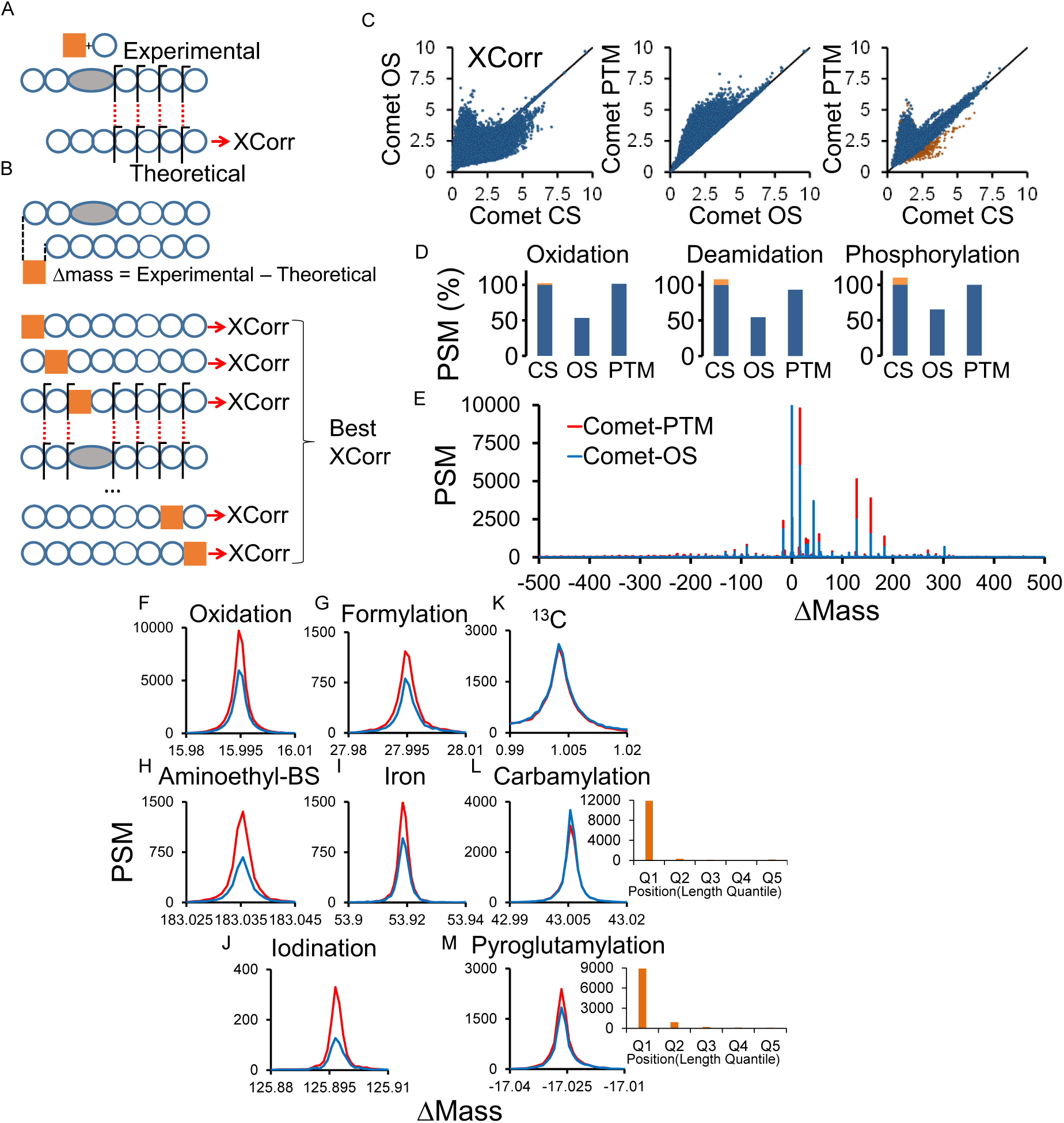
Overview and peptide identification performance of Comet-PTM. (**A**) Conventional OS methods can identify modified peptides from MS/MS spectra, but only the fragments unaffected by the modification (orange square) are matched; this effect diminishes the score assigned to modified peptides, decreasing identification performance. (**B**) Comet-PTM firstly calculates the difference between the mass of the candidate and the mass of the precursor ion detected by the mass spectrometer (ΔMass). ΔMass is then iteratively added to each amino acid in the peptide sequence and the position that yields the best score is selected as the correct match. (**C**) The HeLa dataset from the original OS paper (Chick et al., 2015) was searched using Comet in closed search (CS) mode with 3 variable modifications (Met oxidation, Asn and Gln deamidation and Ser and Thr phosphorylation), with Comet in OS mode, or with Comet-PTM (500 Da tolerance in the later 2 cases). The scores obtained for the same spectra in the different searching conditions are compared. Yellow points in the rightmost graph are PSMs that match peptides with more than one modification in CS mode. (**D**) Identification performance of OS and Comet-PTM relative to CS in the populations of peptides modified by oxidation, deamidation or phosphorylation. The number of PSMs was obtained after filtering by the score threshold corresponding to 1% FDR in the CS. PSMs matching peptides with more than one modification in CS are indicated by orange bars. (**E**) Frequency distribution of PSMs obtained by OS and Comet-PTM as a function of ΔMass. (**F-M**) Details of the frequency distribution around the indicated modifications. The insets at the right in (L) and (M) show the distribution of PSM as a function of the quantile position of the modification in the peptide sequence assigned by Comet-PTM. See also Figure S1.

Here we present a suite of bioinformatic tools designed to overcome these limitations. Our tools double the coverage attained by existing algorithms, so that comprehensive peptide maps can be obtained including practically all the modifications potentially detectable by MS and closed database searches. Our approach also allows accurate location of the modified residues and quantitative analysis of PTMs in the context of proteome-wide studies. We demonstrate the performance of the new tools by performing a comprehensive, tissue-specific characterization of PTMs that are induced by mitochondrial heteroplasmy in a mouse model. Heteroplasmy is a situation that has recently attracted the attention of the biomedical community because it may be produced during mitochondrial replacement therapies aimed to prevent transmission of pathogenic mutations in mitochondrial DNA (Craven et al., 2010) or to treat infertility (Wolf et al., 2015). Mitochondrial heteroplasmy in mice can be genetically unstable and produce adverse physiological effects (Sharpley et al., 2012). The exact molecular mechanisms that produce the pathological effects are unclear and the potential health risk produced by heteroplasmy in the offspring is still a matter under debate. Our results show that heteroplasmy between non-pathological mitochondrial DNA variants induces an array of oxidative modifications in heart that are concentrated in proteins belonging to the oxidative phosphorylation system. The new tools thus improve our ability to interpret the totality of information present in MS/MS datasets and provide new proteome-wide perspectives for systems biology analysis in high throughput proteomics.

## RESULTS

### Comet-PTM enables comprehensive identification of peptide modifications

We developed Comet-PTM, an improved OS engine that takes account of the mass shift produced by the modification in the fragmentation series, producing the same score as a CS search using the same mass increment as a variable modification (Figure 1B). To test the performance of Comet-PTM, the results were compared with those obtained with OS and with CS using 3 common variable modifications. As expected, OS assigned higher scores to a large population of peptide-spectrum matches (PSM) containing modifications not included in the CS list of variable modifications (Figure 1C, left). However, CS assigned a higher score than OS to a large population of PSMs with modifications included in the CS list, because the modifications affected the matching of fragment ions, decreasing the OS-assigned score (Figure 1C, left). This effect reduced the identification performance of OS to around 50% of the PTMs identified by CS (Figure 1D), confirming previous results (Chick et al., 2015). In contrast, scores obtained with Comet-PTM were equal to or better than OS (Figure 1C, center). In addition, Comet-PTM identified the same ΔMass peaks as OS (Figure 1E); however, most of these peaks contained about 2-fold more PSMs (Figure 1E-J), reflecting the superior performance. As expected, this effect was not observed when the modification did not affect the fragmentation series. Thus, the ^13^C peak, produced by errors in assignment of the monoisotopic precursor mass, was observed with exactly the same number of PSMs (Figure 1K). The number of PSMs was also similar for modifications in N-terminal position of the peptide, which only affect b-series and have a negligible effect on identification by HCD fragmentation (Figure 1L,M).

Comet-PTM produced scores similar or higher than those obtained with CS (Figure 1C, right), except for a small population of peptides for which CS found 2 or more modifications (Figure1C, orange dots). This effect was not due to differences between the scores of CS and those of Comet-PTM (Figure S1A and B). Therefore, and unlike OS, the identification performance of Comet-PTM was similar to that of CS for the preselected modifications (Figure 1D). In several instances, Comet-PTM correctly located an oxidation on Trp, Pro or Tyr in the same peptide where CS wrongly assigned the oxidation to Met, the predefined variable modification residue in CS (Figure S1C-E). This finding shows that conventional searches for PTM using a variable modification on selected amino acids can force false assignation of the modified residue. Taken together, these data show that the Comet-PTM OS engine efficiently resolves the modification mass shift in the fragmentation series, doubling the number of modified peptides identified by conventional OS and matching the identification performance of targeted CS.

### Comet-PTM detects the location of modifications in the peptide sequence

Peptides were identified after Comet-PTM searches with SHIFTS, an algorithm that detects the peaks in the ΔMass distribution and controls the peptide false discovery rate (FDR) through a conservative, 3-layered approach (Figure 2A-C; see also Experimental Procedures). From the output of a search against a concatenated target-decoy database, SHIFTS calculates a global score threshold to control FDR in the population of PSMs with ΔMass > −56 Da (Figure 2A). A local score threshold is also defined to control FDR separately within each of the ~1 Da-bins in the ΔMass distribution (Figure 2B). Finally, a peak score threshold is calculated to control FDR separately within each peak detected in the ΔMass distribution (Figure 2C). A PSM is considered correctly identified when its score is above the global and peak thresholds; when a PSM does not form part of a peak, instead of the peak threshold the local threshold is used. This conservative approach allows full control of FDR, avoiding any bias due to the specific behavior of certain kinds of peptide modifications that may be more prone to match decoy sequences.

**Figure 2.**
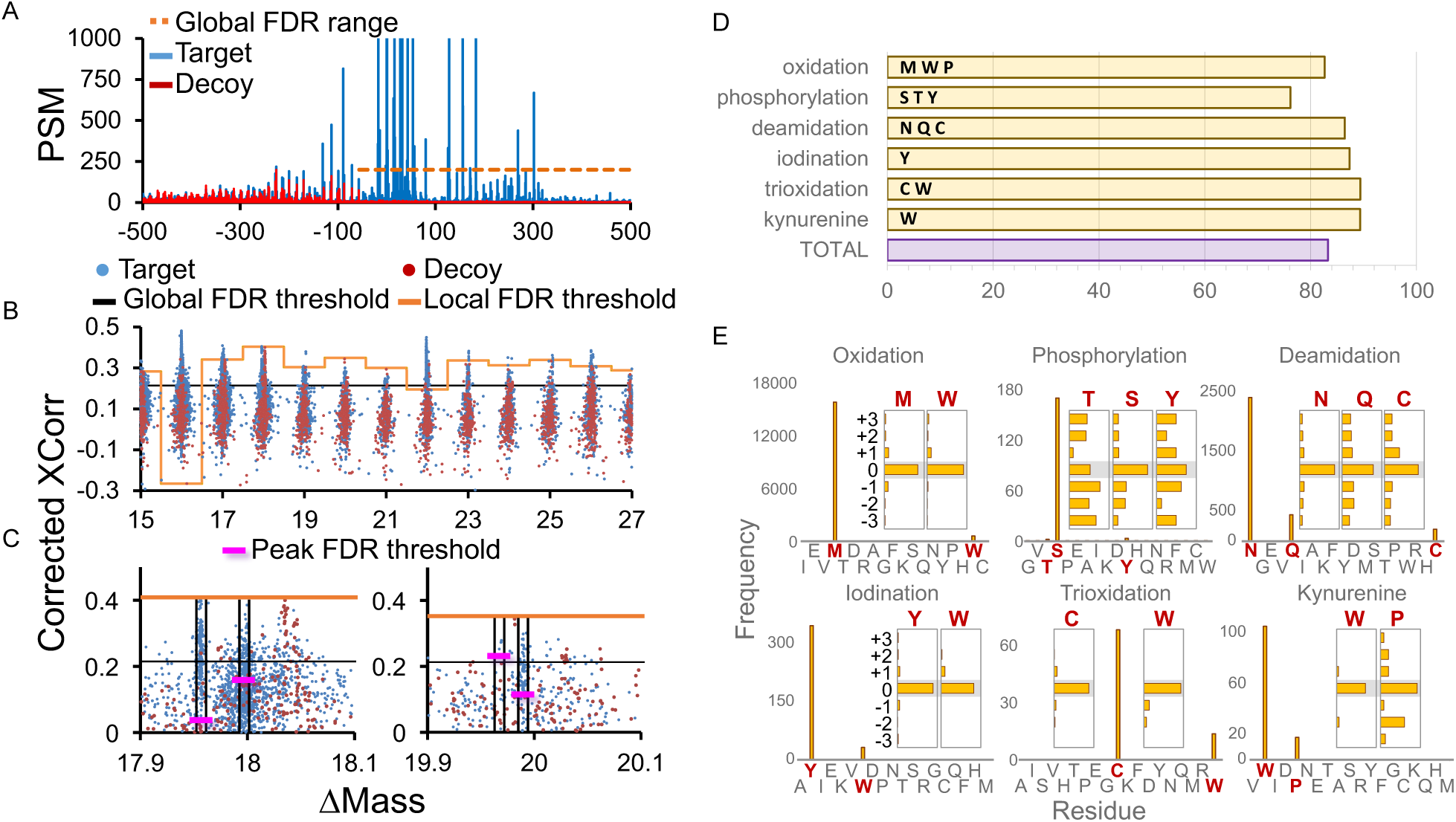
Identification of modified peptides and location of the modified site. (**A**) ΔMass distribution of target and decoy PSMs and range of application of the global score threshold. The vertical scale is enlarged to show the distribution of decoy PSMs. (**B**) Local score thresholds applied in each ~1 Da bin in the ΔMass distribution. (**C**) Peak score thresholds applied to the peaks detected by SHIFTS in the ΔMass distribution. (**D**) Percentage of success assignation of modification sites located in the indicated residues for each one of the displayed modifications. (**E**) Insets: horizontal bar graphs plotting the frequency distributions of PSMs assigned by Comet-PTM to the indicated amino acids and modifications. Vertical bar graphs: frequency of PSMs assigned to the indicated amino acids after substracting the average frequencey of the three previous and the three subsequent positions. Residues that accumulate counts in the histogram at position 0 are labelled in red. See also Figure S2.

An advantage of Comet-PTM over existing OS approaches is that it automatically assigns the modification to the residue in the peptide sequence that produces the best score and therefore is the modified position that best explains the fragmentation data. Assigning modifications to specific residues is considered a much less reliable process than identifying peptides, partly because there is frequently insufficient information to determine the exact modified residue (Chalkley et al., 2008). To estimate the accuracy of Comet-PTM assignation of a modification to the correct site, we filtered correct identifications using the conservative method described above using 1% FDR threshold and then calculated the fraction of PSMs in which the mass shift was located in amino acids predicted to harbor well-known modifications. Oxidation was located to Met, Trp or Pro in 82% of cases, and deamidation to Asn, Gln or carbamidomethylated Cys in 86% of cases (Figure 2 D). Considering six well-known modifications, the overall accuracy of Comet-PTM was estimated to be around 85%.

We then analyzed which amino acids were assigned to known modifications with a frequency above that expected by chance. Among the most frequent, we only found well-known modifications like oxidized Met and Trp, deamidated Asn, Gln and carbamidomethylated Cys, phosphorylated Ser, trioxidized Cys and Trp and kynurenine-modified Trp (Figure 2E). Some Pro modifications were also found with the ΔMass of kynurenine-modified Trp (Figure 2E), but all of them corresponded to homologous peptides that contained the Pro > Thr substitution, which has the same ΔMass (Figure S2), and that were assigned the same score. Finally, Trp iodination was also clearly detected above background (Figure 2E). This modification was unexpected because although Trp halogenases have been described in bacteria (van Pee and Patallo, 2006) they are not present in mammals, and Trp iodination has not been described before as a chemical artifact. The majority of peptides containing iodinated Trp contained neither Tyr nor His, the 2 amino acids known to react with iodine. A careful inspection of their fragment spectra confirmed the presence of y-fragments that could only be explained by Trp being the modified residue (Figure S3). In addition, the only modification described in Unimod that matched the observed mass shift corresponded to iodination. From all these data we concluded that Trp was iodinated in these peptides, most probably as a side-chain reaction of the iodine produced from the Cys-alkylating reagent iodoacetamide. This finding further highlights the accuracy of Comet-PTM to locate the site of modification and suggests that this OS engine may be useful for detecting novel peptide modifications in specific amino acids.

### A single integrated statistical framework allows quantification of the proteome and of the modified peptidome

For the quantitative analysis of modified peptides, we developed a novel algorithm based on a previously proposed systematic workflow (Garcia-Marques et al., 2016; Navarro et al., 2014) (Figure 3A). The algorithm includes a peptide-to-protein integration step that quantifies protein values from the unmodified peptide forms and then computes the standardized log2-ratio of the modified peptides with respect to these protein values (*Z*_*pq*_) (Figure 3B). This makes it possible to detect modified peptides whose behaviors deviate significantly from those of the unmodified peptides from the same protein. Note that the algorithm calculates peptide abundances relative to the parent protein, not to the mean of all the peptides in the sample, so that changes at the peptide level are unaffected by changes in protein abundance. When applied to a biological model, the *Z*_*pq*_ distributions of the unmodified and modified peptide forms precisely followed the expected null hypothesis distribution in different tissues from 2 mouse strains (Figure 3C). These results convincingly demonstrate that the quantitative behavior of the entire population of modified peptides can be modelled accurately, providing a robust statistical framework within which to analyze abundance changes in high-throughput experiments. This allowed detection of significant changes in certain modified peptides against the null hypothesis in specific situations (Figure 3C, lower left).

**Figure 3.**
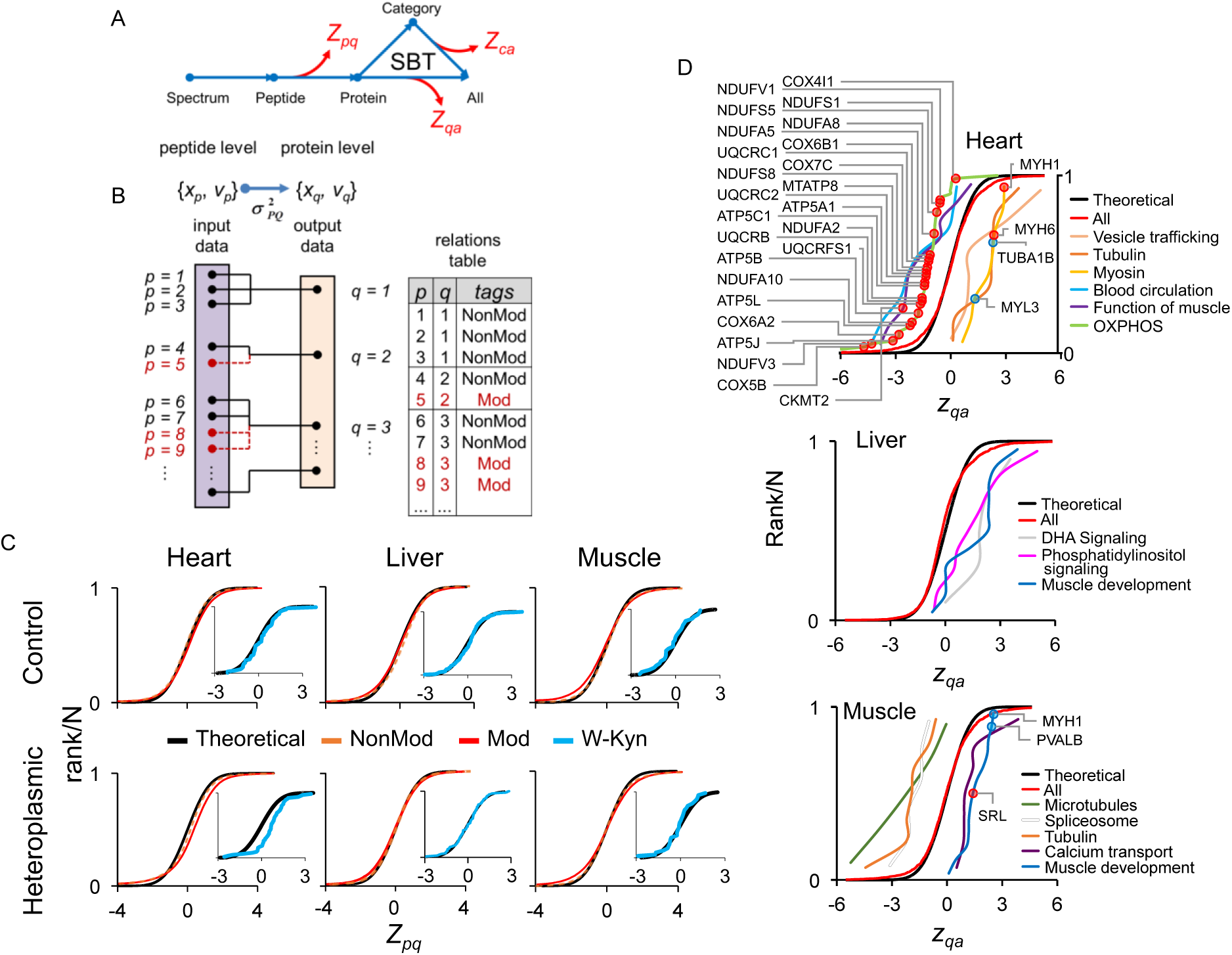
Quantitative analysis of alterations produced by heteroplasmy in the modified peptidome and in the proteome. (**A**) Scheme of the integrative workflow used to quantify modified peptides, proteins or categories. Each arrow represents a step performed with the generic integration algorithm (Garcia-Marques et al., 2016). Standardized log_2_-ratio values at the peptide level (*Z*_*pq*_) are obtained from the peptide-to-protein integration. The algorithm provides corrected peptide values by taking account of the corresponding protein changes. The same workflow is used for protein quantification and systems biology analysis of coordinated protein responses with the SBT model (Garcia-Marques et al., 2016).**(B)** Scheme of the Generic Integration Algorithm (GIA) (Garcia-Marques et al., 2016) adapted for the analysis of modified peptides in the peptide-to-protein integration step. The relations table include tags that can be used to indicate that modified peptides are excluded from the computation of protein averages.(**C**) Distributions of *Z*_*pq*_ values for non-modified (Non-Mod) and modified peptides (Mod). These data come from the analysis of mouse tissues in a model of mitochondrial heteroplasmy; each graph correspond to a tissue from an individual animal. The inset plots show the *Z*_*pq*_ distributions for peptides containing the kynurenin Trp modification. Note that the distribution is displaced to the right in the hearts of heteroplasmic mice, indicating increased abundance. For further details see Experimental Procedures.(**D**) Characterization of coordinated protein alterations produced by heteroplasmy in heart, liver and muscle using the SBT model. *Z*_*qa*_ values are standardized log2-ratio averages of proteins from heteroplasmic mice relative to controls. Proteins are grouped into representative functional categories with a statistically significant change. The upper panel presents proteins in the OXPHOS category that are decreased in the heteroplasmic heart and that contain heteroplasmy-induced oxidative modifications (Figure 6). See also Table S1.

The same workflow used for peptide quantitation allowed the analysis of protein abundance changes and the characterization of functional categories that were affected by the coordinated action of proteins, using the Systems Biology Triangle (SBT) model (Garcia-Marques et al., 2016) (Figure 3A). This statistical design thus allowed a full description of the alterations in the modified peptidome in the global context of changes in the proteome.

### Heteroplasmy produces protein alterations consistent with a mitochondrial dysfunction in heart

We used Comet-PTM and associated tools to study the molecular impact of mitochondrial heteroplasmy on the proteome and the modified peptidome in heart, liver and skeletal muscle from 12-week old mice, in a model we describe in detail in another report (Latorre-Pellicer et al., submitted). SBT model analysis of the quantitative protein data from the heart revealed a coordinated decrease of proteins related to muscle function and the oxidative phosphorylation system (OXPHOS) and a coordinated increase of tubulins, myosins and proteins involved in vesicle trafficking. The muscle-function proteins included KCRS (mitochondrial creatine kinase) and VDAC3, which are key suppliers of phosphocreatine to the sarcomere, and SGCG and KGP1, which are essential for sarcomere contraction (Table S1). These alterations found in heteroplasmic mice are highly consistent with the decreased ATP synthesis and the abnormal increase in phosphocreatine/ATP ratio in heart and with the increase in plasma creatine kinase we have reported in this animal model (Latorre-Pellicer et al., submitted), and provide evidence that heteroplasmic cardiac mitochondria have a compromised ability to supply energy to the sarcomere. Among the increased vesicle trafficking proteins, STXB3 and VAMP2 are implicated in the insulin-dependent movement of GLUT4 from inner vesicles to the plasma membrane. This finding also agrees with the higher glucose uptake we detected in heteroplasmic hearts (Latorre-Pellicer et al., submitted), pointing to a shift toward glycolytic metabolism in order to compensate the reduced phosphocreatine supply from OXPHOS. The coordinated decrease in blood proteins and the increase in heart cytoskeletal proteins in heteroplasmic animals suggest a homeostatic effort to maintain cardiac cell structure.

Heteroplasmy had not effect on OXPHOS protein levels in liver, but did produce a coordinated increase of DHA signaling pathway proteins (Figure 3D), including AKT2, BID and B2CL1 (Table S1). AKT2 plays an essential role in the progression of the inflammatory response and apoptosis (Lopez-Carballo et al., 2002); consistent with this finding, heteroplasmic mice show signs of inflammation in the liver with aging (Latorre-Pellicer et al., submitted). Heteroplasmic liver also showed coordinated increases in a group of myosins and phosphatidylinositol pathway proteins, probably reflecting a homeostatic response. These data thus indicate that at the age of 12 weeks heteroplasmy produces a decrease in OXPHOS proteins in heart, probably reflecting mitochondrial dysfunction, but not in liver.

In heteroplasmic muscle we detected a coordinated decrease of proteins regulating cell shape, such as tubulins and microtubule proteins (Figure 3D). On the other hand, the thick-filament myosins, as well as calcium transporting proteins, were increased, suggesting the induction of compensatory mechanisms. However, we found no evidence of altered expressión of mitochondrial proteins in muscle at this age.

### Heteroplasmy induces oxidative modifications of OXPHOS proteins in heart

To study the impact of heteroplasmy on the modified peptidome, we first compared the ΔMass distribution of peptides searched with Comet-PTM and identified with SHIFTS in the 3 tissues. The distribution of the most abundant peaks was very similar in liver, heart and muscle (Figure S3). In addition to the most intense peak corresponding to unmodified peptides, we found peaks corresponding to oxidized and deamidated peptides, and dioxidized peptides were also frequent. Other frequent modifications were artifacts produced by TMT reagents and iodoacetamide treatments, sodium adducts and missed cleavages. Some tissue-specific modifications were also detected, mostly in the region from 50 to 100 Da. Among these, phosphorylated peptides were more abundant in muscle and 90% of these peptides belonged to proteins implicated in muscle contractility, consistent with the regulatory role of phosphorylation in the sarcomere. Within each Δmass peak, we analyzed the distribution of the modified amino acid residues which were assigned to frequent modifications using the unbiased approach followed in Figure 2E. Confirming the results described above, amino acids known to harbor specific modifications were mostly located correctly (Figure S4). Interestingly, the relative proportion of modified sites was remarkably conserved across the 3 tissues, with the exception of tyrosine iodination, which was prominent in heart but less frequent in liver and skeletal muscle (Figure S4). This analysis also detected the increased frequency of phosphorylation in muscle (Figure S4). We estimated that about one quarter of the total peptidome in the 3 tissues had known posttranslational modifications, mostly oxidation and deamidation; less than 10% of modifications were chemical artifacts (Figure 4A).

**Figure 4.**
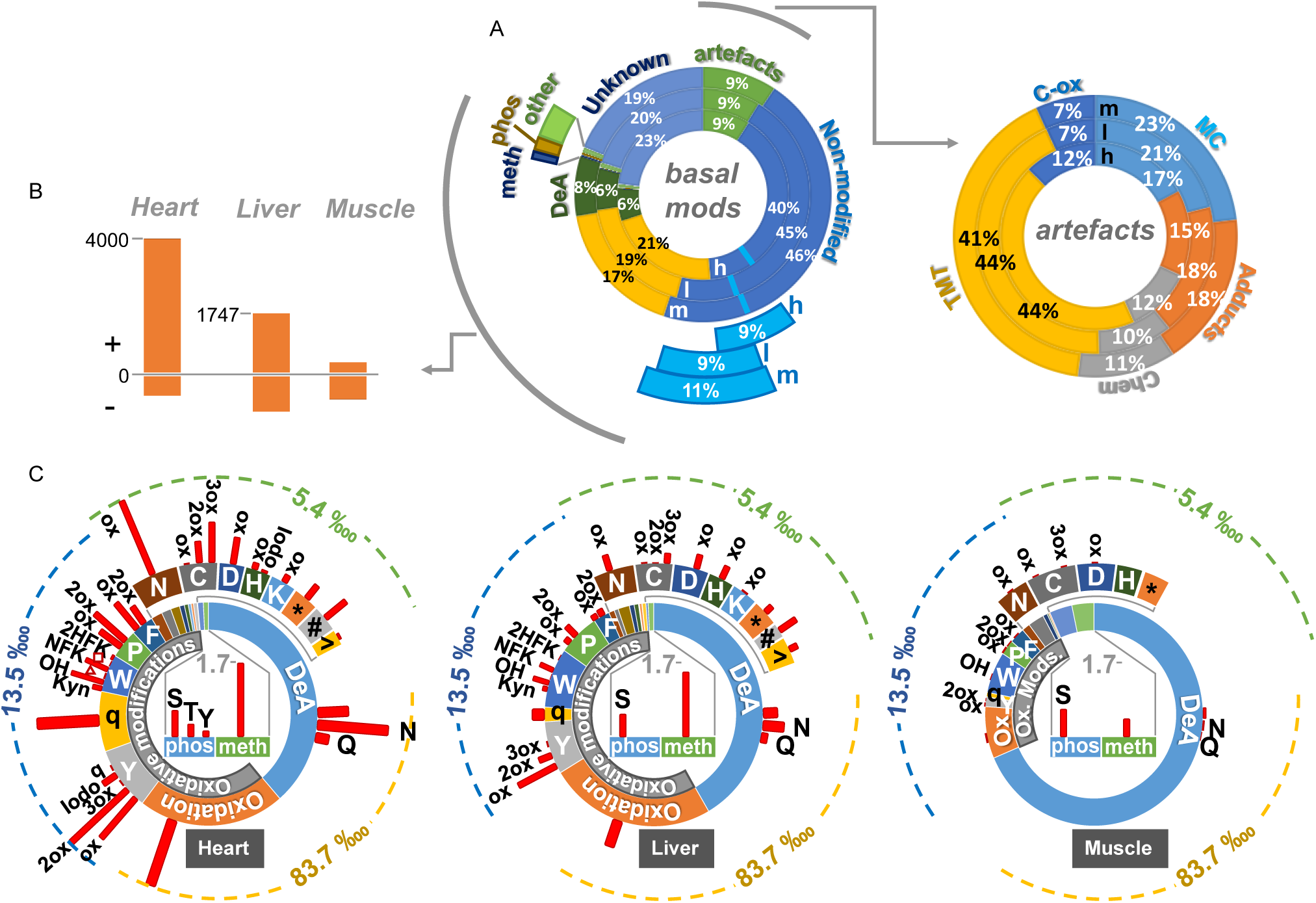
Alterations produced by heteroplasmy in the modified peptidome in different mouse tissues. (**A**) Complete map of the basal peptidome, expressed as a percentage of the total number of peptides identified in each tissue (circular bar graphs: h: heart; l: liver; m: muscle). For simplicity, peptides oxidized on Met (light blue) are included in the group of non-modified peptides. The plot to the right shows proportions of artifacts produced by sample manipulation. (**B**) Number of PTM-containing peptides significantly increased (+) or decreased (-) in heteroplasmic mouse tissues (n=4) relative to controls (n=3). (**C**) Statistically significant PTM increases in heteroplasmic mouse tissues according to the type of modification and the modified residue. The circular inner bars show the peptide proportion for each modification in the basal state, and the radial red bars represent the proportion of peptides of each kind (in parts per ten thousand) that are increased in heteroplasmic tissues. For the shake of clarity, 3 different scales are used depending on the type of modification. DeA: deamidation; meth: methylation; phos: phosphorylation; MC: missed cleavages; Chem: chemical derivatives produced by sample preparation; TMT: extra addition of the isobaric labeling reagent; Adducts: Na, K and ammonia adducts. Capital letters indicate the single letter aminoacid code. ox: oxidation; 2ox: dioxidation; 3ox: trioxidation; Kyn: Trp to kynurenine; OH: Trp to hydroxytryptophan; NFK: Trp to N-formyl kynurenine; 2HFK: Trp to 2-hydroxy formyl kynurenine; red triangle: Trp to oxolactone; red square: Trp to quinone; q: quinone, Tyr to quinone; Iodo: iodination; *: Met to homocysteic acid; #: Met to homoserine. See also Figures S3 and S4 and Table S2.

Secondly, we studied whether heteroplasmy altered the pattern of modifications in each tissue by analyzing the quantitative data with the integrative statistical algorithm (Figure 3A). Heteroplasmy induced marked tissue-specific modifications, most of them concentrated in heart and barely detectable in skeletal muscle (Figure 4B). The vast majority of increased modifications in heart were oxidative, affecting mostly to Tyr, Trp, Pro and Phe and to a lesser extent Asn, Cys and Asp (Figure 4C). Heteroplasmy also induced phosphorylation and methylation in the heart. The pattern of modifications was similar in liver, but the effect of heteroplasmy on this tissue was significantly less pronounced.

Functional enrichment analysis revealed that heteroplasmy-induced modifications in mice heart were mostly located in mitochondrial proteins, particularly those located in the inner mitochondrial membrane; moreover, most of the modified proteins were OXPHOS proteins, which were enriched with multiple oxidative modifications (Figure 5A). TCA cycle proteins and those related to the oxidoreductase complex were enriched in several modifications. Deamidation affected several mitochondria-related categories, and phosphorylation affected proteins related to contractility. Heteroplasmy-induced modifications affected considerably fewer categories in liver (Figure 5B) and were not significantly enriched in muscle (Figure 5C), where only a few decreased categories were detected, containing low numbers of proteins. Deamidated peptides were found in proteins related to mitochondria, NAD binding and extracellular exosomea in both heart and liver, suggesting that these modifications do not participate in the tissue-specific phenotypic differences between heteroplasmic and control mice (Latorre-Pellicer et al., submitted).

**Figure 5.**
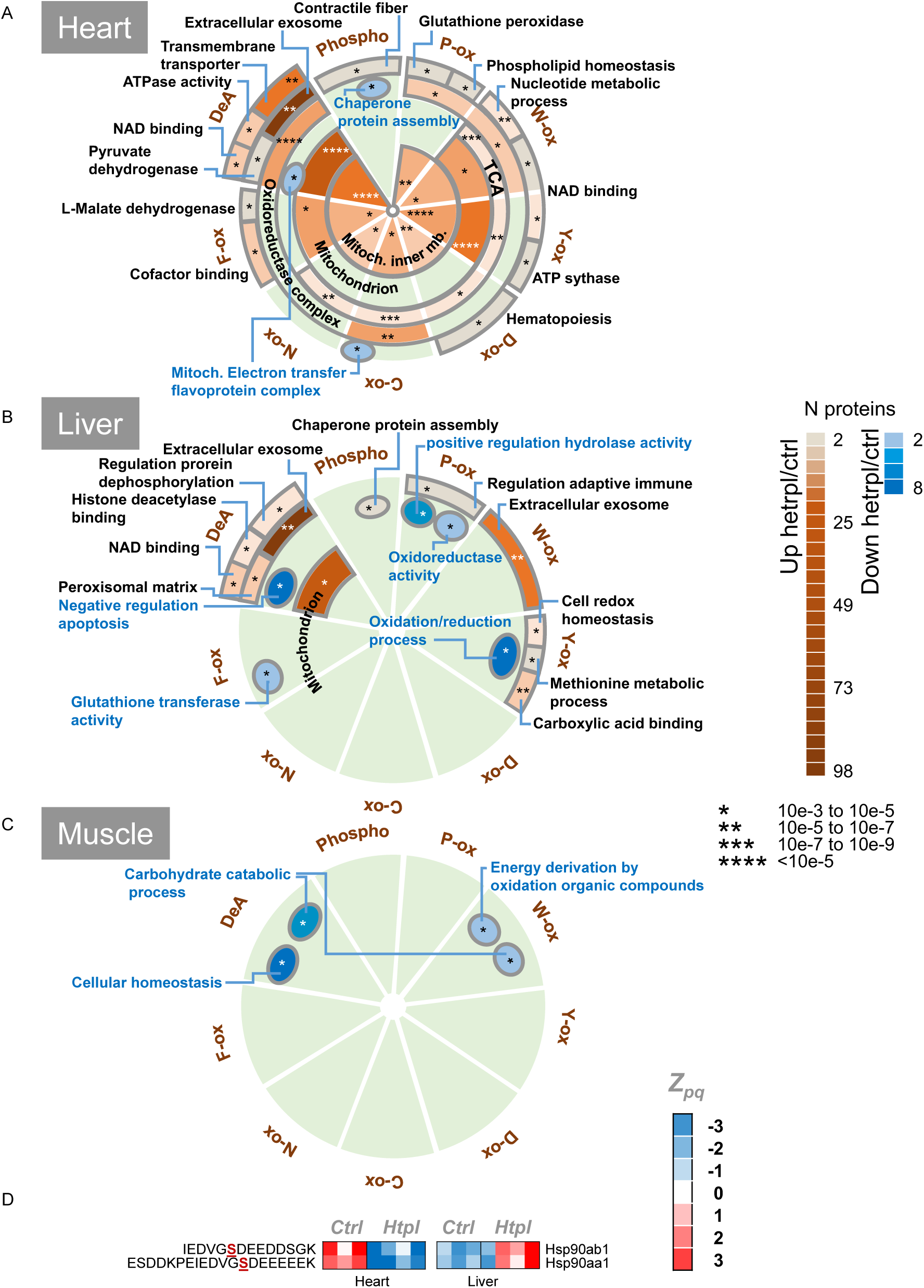
Function distribution of proteins posttranslationally modified by heteroplasmy. (**A-C**) Functional enrichment analysis of the proteins containing PTMs that are significantly increased (brown scale) or decreased (blue scale) in different tissues of heteroplasmic mice in relation to control mice. The analysis was performed separately per different kinds of modification (slices). Color intensity represents the number of proteins in each category. Asterisks indicate statistical significance of category enrichment relative to the list of proteins identified in each tissue. Inner circular rings are used for categories at the cellular component level, whereas outer rings correspond to biological processes. (**D**) Heat map of the *Z*_*pq*_ values for 2 modified proteotypic peptides from the 2 HSP90 isoforms (HSP90AB1 and HSP90AA1) for all control and heteroplasmic replicates in the heart and in liver. These 2 isoforms are included in the “chaperone protein assembly” functional category displayed in **A** and **B**.

Interestingly, heteroplasmy decreased phosphorylations in proteins related to chaperone activity in the heart but had the opposite effect in liver (Figure 5A and B). This category contains the two HSP90 isoforms (HSP90AA1 and HSP90AB1), which harbor PTM sites involved in the regulation of the chaperone activity (Mollapour and Neckers, 2012). These isoforms are known to be phosphorylated at Ser263 and Ser255, respectively (Hu et al., 2015; Wang et al., 2017), within the CK2 phosphorylation motif (S-X-X-E) that is crucial for protein activity. In particular, phosphorylation at Ser255 is required for the activation of MAPK/ERK pathway. Comet-PTM and SHIFTS identified these 2 phosphorylated sites, and heteroplasmy decreased their levels in heart while increasing them in liver (Figure5D), suggesting that heteroplasmy differentially affects chaperone activity. These data support the role we have proposed for the endoplasmatic reticulum and the mitochondrial unfolded stress response in cardiac heteroplasmy (Latorre-Pellicer et al., submitted).

To compare the alterations in the modified peptidome with the changes observed at the protein level, we mapped the proteins that harbored post-translational modifications affected by heteroplasmy into the protein distribution plots that depicted coordinated protein responses (red and blue dots in Figure 3D). The majority of increased modifications were mapped into the OXPHOS protein category in heart, which was decreased, while the proteins belonging to the rest of categories were mostly unaffected. This finding suggests that the oxidative damage produced by heteroplasmy in heart OXPHOS proteins induced their degradation, and is in good agreement with the increase in oxidative stress observed in heart slices, embryonic fibroblasts and adult fibroblasts of heteroplasmic mice (Latorre-Pellicer et al., submitted).

Among the OXPHOS proteins affected by heteroplasmy, most oxidative modifications occurred in the 4 respiratory protein complexes, except for Trp oxidations, which were mostly concentrated in C-IV (Figure 6A). The modifications were clustered in groups of proteins located together in the structure of the complexes, and predominantly affected proteins orientated toward the mitochondrial matrix (Figure 6), where proteins are more exposed to mitochondrial ROS. In further analysis, we used Consurf to calculate evolutionary conservation profiles for the affected residues (Ashkenazy et al., 2016) and mapped the modifications onto the known 3D structures of the complexes, when available, or onto their predicted models (Meier and Soding, 2015). We found that 70% of the modified residues were conserved or interacted with conserved regions (Table S3). Moreover, most of the modifications were located on the protein surface, and those that were buried tended to have high conservation scores, suggesting they are important for structural stability or folding (Table S3). These findings suggest that these oxidative modifications compromise OXPHOS protein function. This viewpoint is supported by the decreased ATP synthesis capacity we have determined in heteroplasmic heart (Latorre-Pellicer et al., submitted).

**Figure 6.**
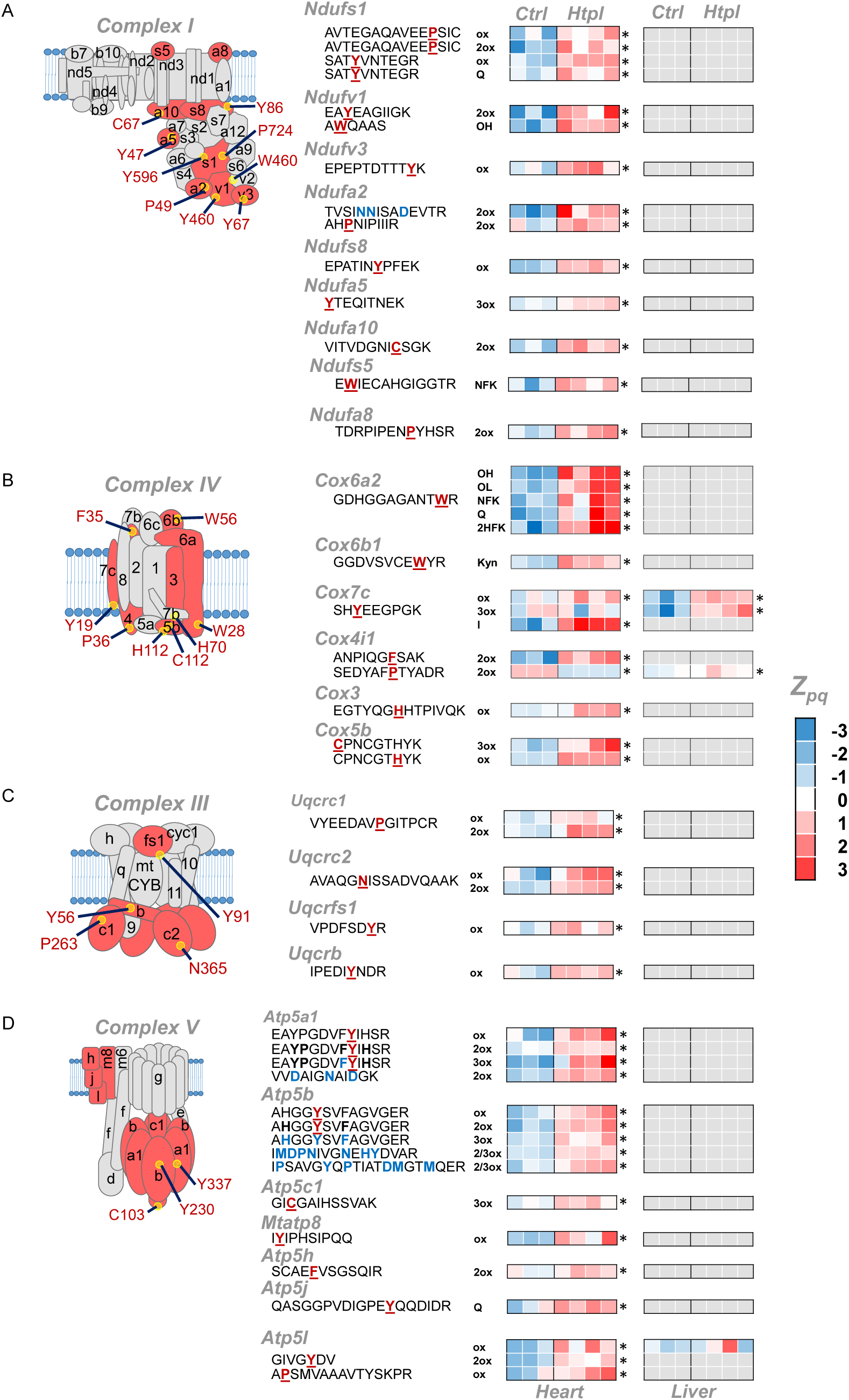
Posttranslational modifications induced by heteroplasmy in OXPHOS proteins. (**A-D**) Scheme of heteroplasmy-induced PTMs in proteins of complex I, complex III, complex IV and complex V from the OXPHOS system. PTM-containing proteins are highlighted in red, with the spatial position of the modified residue indicated. The heat maps show statistically significant changes in abundance (*Z*_*pq*_) of the indicated peptides in heart and liver from control mice (Ctrl) and heteroplastic mice (Htpl). Grey squares indicate that the modified peptide was not detected in the tissue sample. Modified residues that are unambiguously assigned are indicated with underlined bold red letters; tentatively assigned sites are labelled in bold blue letters. Asterisks mark statistically significant changes between control and heteroplasmic samples according to Student’s *t*-test (p < 0.05). The modification type is indicated in the column to the right of the peptide sequences: ox: oxidation; 2ox: dioxidation; 3ox: trioxidation; Kyn: Trp to kynurenine; OH: Trp to hydroxytryptophan; NFK: Trp to N-formyl kynurenine; 2HFK: Trp to 2-hydroxy formyl kynurenine; OL: Trp to oxolactone; Q: Trp to quinone; I: iodination. The conservation scores of these peptides are listed in Table S3, from which the results of Vseq analysis can also be downloaded. See also Table S3.

## DISCUSSION

Comprehensive analysis of PTMs in mass spectrometry data obtained through shotgun approaches has historically presented a bioinformatics challenge because conventional (or closed) database searches can only identify predefined subsets of modifications. This limitation impedes truly hypothesis-free analysis of modifications, to the point that the proportion of chemical and posttranslational modifications of proteins in biological systems has remained unknown. The open database search strategy (Chick et al., 2015) provided a starting point for resolving this problem, allowing a truly unbiased high-throughput identification of modified peptides without previous knowledge on the nature of these modifications. An improved fragment-ion indexing method (MSFragger) later demonstrated that open searches can be performed on a timescale similar to or shorter than that required for closed searches (Kong et al., 2017). The open search strategy has the additional advantage that modified and non-modified peptide forms are identified simultaneously (Kong et al., 2017). These strategies, however, are not comprehensive and provide limited information on the nature of modified residues. The computational approach presented here enables the open search strategy to match the performance of closed searches, identifying the vast majority of detectable peptide modifications. Although Comet-PTM is built on the framework of a well-known database searching engine, it is important to note that the underlying concept is extrapolable to any other searching engine, opening the way to near-complete analysis of the protein-modification landscape in biological systems using tools already used by the proteomics community. In addition, our method provides information on the most probable site containing the modification with an accuracy estimated to be around 85%. We should note that, in its current state of development, the method might fail the peptide contains more than one modification. However, the impact of this limitation on the general performance of the algorithm seems small, since the fraction of multiply modified peptides that were not identified was almost negligible. We also introduced a conservative, three-layered approach to control the FDR of peptide identification. Our local FDR calculation is similar to that used by MSFragger (Kong et al., 2017), with the only difference that we used the decoy/target competition strategy to estimate FDR, while Kong et al. modeled local score distributions, in the same 1 Da bins, as a mixture of correct and incorrect identifications and calculated the probability of correct identification by the Bayes rule. As recognized by these authors, their approach did not take into account the p.p.m. levels of accuracy produced by high-resolution instruments (Kong et al., 2017), which we have considered in our model by applying an additional peak FDR estimation. In addition, we found it necessary to introduce a third global FDR filter to avoid identification of low-scoring peptides in local bins or in peaks that, as in the case of oxidation, concentrate high numbers of target PSMs.

The new algorithm allowed us to produce peptide maps containing realistic proportions of unmodified peptides, posttranslational modifications, and chemical artifacts in mouse tissues, and of amino acid residues subjected to each type of modification. We show that the basal distribution of modifications and modified amino acids is remarkably similar across the 3 mouse tissues studied, indicating that the algorithm generates reproducible results and that most peptide modifications are of an oxidative nature. Many of the modifications are introduced during sample preparation, including those induced by exposure to Cys-alkylating and amine-directed isobaric labeling reagents; while these agents are thought to be highly specific, in the practice they produce secondary reactions with other nucleophiles.

The ability to perform a truly hypothesis-free high-throughput analysis of modifications can contribute to a deeper understanding of the relation between protein chemistry and function; however, the most important biological insight obtainable with shotgun proteomics is how these modifications are altered during normal physiological processes or as a consequence of disease. The statistical approach presented here enables quantitative analysis of the modified peptidome within the same framework used for the analysis of unmodified proteome; this approach thus allows coherent interpretation of all the results and opens the way to integrated systems-biology analysis. We demonstrate the performance of the new algorithm by simultaneously characterizing, in 3 tissues of a mouse model, the quantitative impact of heteroplasmy on the proteome and on the modified peptidome. To our knowledge, this is the first study to apply open-search-based approaches together with quantitative mass spectrometry, allowing comprehensive and unbiased characterization of alterations in the protein modification landscape produced in a biological system.

Our results reveal that the major hallmark of mitochondrial heteroplasmy in 12-week old animals is oxidative damage to OXPHOS proteins in the heart. This is reflected in the increased levels of oxidative modifications and in the decreased levels of OXPHOS proteins. Our data support the notion that heteroplasmy between different non-pathological mtDNA variants affects the performance of the OXPHOS system in the heart, resulting in a compromised mitochondrial ROS handling that triggers an oxidative stress response and impairs the ability to supply energy to the sarcomere. Heteroplasmy is known to be related to ROS production (Hämäläinen et al., 2015; 2013) and also affects the levels of contractile proteins in muscle. These alterations do not occur in liver, which is able to resolve heteroplasmy (Latorre-Pellicer et al., submitted). Our data thus provide a molecular mechanism that explains the functional findings reported in heteroplastic heart, including a diminished ability of mitochondria to produce ATP, increased glucose uptake, an exacerbated phosphocreatine/ATP ratio, and mitochondrial disruption and structural alterations (Latorre-Pellicer et al., submitted).

## AUTHOR CONTRIBUTIONS

N.B., E.B., E.C. and J.V. designed the algorithms. S.M. and M.T-H. programmed Comet-PTM code. N.B. programed SHIFTS and analyzed Comet-PTM performance. N.B., E.B. and M.T-H. programmed the new generic integration algorithm and performed the quantitative analysis of PTM. J.A.E., A.V.L-V. and A.L-P. developed the heteroplasmic mouse model. A.V.L-V., R.M. and E.C. prepared the tissue extracts. R.M. and E.C performed the mass-spectrometry analysis. I.E. and J.M.R. performed the bioinformatic analysis of PTM. N.B., E.B., E.C. and J.V. wrote the manuscript. J.V. directed the research.

## ACKNOWLEDGEMENTS

We thank Simon Bartlett (CNIC) for English editing. This study was supported by competitive grants from the Spanish Ministry of Economy and Competitiveness (MINECO) (BIO2015-67580-P) through the Carlos III Institute of Health-Fondo de Investigación Sanitaria (PRB2, IPT13/0001 - ISCIII-SGEFI/FEDER, ProteoRed), the Fundación La Marato TV3 and by the FP7-PEOPLE-2013-ITN *Next generation training in cardiovascular research and innovation-Cardionext*. Navratan Bagwan is a FP7-PEOPLE-2013-ITN-Cardionext fellow. The CNIC is supported by the MINECO and the Pro-CNIC Foundation, and is a Severo Ochoa Center of Excellence (MINECO award SEV-2015-0505).

## EXPERIMENTAL PROCEDURES

### Benchmarking mass spectrometry data set

To test the performance of the developed algorithms, we used the publicly available HEK293 dataset (Chick et al., 2015), containing 1.121.149 MS/MS spectra in 24 raw files acquired on a Q-Exactive Orbitrap mass spectrometer.

### Mouse model of heteroplasmy

The mice used in this work were 12-week-old control (C57BL/6JOlaHsd strain) and heteroplasmic mice (containing more than one mtDNA in the same cytoplasm, C57BL/6 background); further details are described in the accompanying manuscript (Latorre-Pellicer et al., submitted). The effect of heteroplasmy on the post-translational modifications of the proteome of different tissues was studied on liver, heart and skeletal muscle (gastrocnemius) samples. In the last two tissues, the heteroplasmy was stable, while the liver was selected as a control tissue since it spontaneously selected one of the alternative variants of mtDNA (Latorre-Pellicer et al., submitted). For each tissue, biological replicates from different control (N=3) and heteroplasmic mice (N=4) were analyzed.

### Preparation of protein extracts

Mice were sacrificed by cervical dislocation and liver, heart and skeletal muscle tissues were extracted. 20 mg of each tissue were homogenized in lysis buffer (10mM Tris-HCL pH7.4, 1 mM EDTA, 0.32 M sucrose, 2% SDS) freshly supplemented with protease and phosphatase inhibitors (Roche) and 50 mM DTT, using a MagNA Lyser instrument (Roche). The lysate was boiled for 5 min and cell debris were removed by centrifugation.

### Protein digestion, peptide labeling and fractionation

Proteins were treated with 50 mM iodoacetamide (IAM) and digested with trypsin using the Filter Aided Sample Preparation (FASP) digestion kit (Expedeon) (Wisniewski et al., 2011) according to manufacturer’s instructions. Dried peptides were labeled using 10 plex-TMT reagents according to manufacturer’s instructions (Thermo Fisher Scientific), desalted on OASIS HLB extraction cartidges (Waters Corp.) (Leyfer and Weng, 2005), separated into 7 fractions using the high pH reversed-phase peptide fractionation kit (Thermo Fisher Scientific) and dried-down before MS analysis.

### LC-MS analysis

Each fraction of the labeled peptide samples were analyzed using an Easy nano-flow HPLC system (Thermo Fisher Scientific) coupled via a nanoelectrospray ion source (Thermo Fisher Scientific, Bremen, Germany) to a Q Exactive HF mass spectrometer (Thermo Fisher Scientific, Bremen, Germany). C18-based reverse phase separation was used with a 2-cm trap column and a 50-cm analytical column (EASY column, Thermo). Peptides were loaded in buffer A (0.1% formic acid (v/v)) and eluted with a 240 min linear gradient of buffer B (80% acetonitrile, 0.1% formic acid (v/v)) at 200 nL/min. Mass spectra were acquired in a data-dependent manner, with an automatic switch between MS and MS/MS using a top 15 method. MS spectra were acquired in the Orbitrap analyzer with a mass range of 400–1500 m/z and 60,000 resolution. HCD fragmentation was performed at 27 of normalized collision energy and MS/MS spectra were analyzed at 60,000 resolution in the Orbitrap.

### Database search

Unless indicated otherwise, all searches were performed using Comet release 2016.01 (Eng et al., 2015; Eng et al., 2013) using trypsin digestion with 1 missed cleavages (unless otherwise specified) and fixed Cys carbamidomethylation (57.021464 Da). For heteroplasmic mice data, TMT labeling at N-terminal end and Lys was also considered as a fixed modification (229.162932 Da). Fragment ion tolerance was 0.02 bin, 0 mass offset. Precursor tolerance type and isotope error were set to 1. Precursor charge range was 2-4, maximum precursor charge 5 and maximum fragment charge 3. Only y- and b-ions were used for scoring.

Closed searches (CS) were performed at 5 ppm precursor ion tolerance, using three dynamic modifications: Met oxidation (15.994915), Asn and Gln deamidation (0.984016) and Ser and Thr phosphorylation (79.966331). Peptide identification from MS/MS data was performed using the probability ratio method (Martinez-Bartolome et al., 2008). False discovery rates (FDR) of peptide identifications were calculated using the refined method (Bonzon-Kulichenko et al., 2015; Navarro and Vazquez, 2009); 1% FDR was used as the default criterion for peptide identification.

Open searches (OS) with Comet and Comet-PTM were performed in the same conditions as CS, except that precursor ion tolerance was set to 500 Da.

### Comet-PTM

Comet.PTM was developed by modifying the open-source database search engine (Eng et al., 2015). For every sequence candidate Comet-PTM calculates the difference between theoretical and experimental precursor mass (ΔMass), and adds up this mass iteratively to each one of the amino acid masses in the peptide sequence, calculating a Xcorr score in each one of the possible modified forms of the peptide (Figure 1B). The selected candidate is the modified peptide form that produces the highest Xcorr. This design allows Comet-PTM to reach the score that would have been obtained by performing a targeted CS with the same modification in the same position. Note that the scores are not exactly identical, since CS uses the theoretical mass of the modification and Comet-PTM estimates it from the difference between the precursor mass and the theoretical mass of the non-modified peptide, and experimental errors on this estimate may affect fragment matching. This effect is, however, small when low ppm precursor mass accuracies are used (Figure S1A and B). Comet-PTM has a user-selectable option of scoring also the non-modified peptide sequence (even when ΔMass is different from zero), to take into account labile modifications (Kong et al., 2017).

Comet-PTM was developed to take full advantage of the multi-thread design of Comet. Comet-PTM used less than 4 hours to perform a 500 Da-wide open search of 16 LC-MS runs, containing an average of 44.390 MS/MS spectra each, using a computer cluster with 16 nodes, where each node is built of 2 x Intel Xeon E5-2695v2 at 2.40 GHz and contained 46 threads/124 gigabyte.

### SHIFTS

SHIFTS (Systematic Hypothesis-free Identification of modifications with controlled FDR based on ultra-Tolerant database Search) is a program that identifies peaks in the ΔMass distribution, assigns PSM to peaks and calculates FDR for peptide identification (Figure S6). SHIFTS uses as input the Thermo .raw files and the files obtained from Comet-PTM search.

#### Mass recalibration

SHIFTS firstly recalibrates precursor peptide masses independently in each raw file. This was done by selecting a population of non-modified peptides with a very high score (user selectable; recommended values are those yielding 0.1% global FDR or lower), which are assumed to be true identifications and are used to calculate the systematic mass error (median deviation in m/z scale), which is assumed to be constant in each raw file. From these data, SHIFTS also calculates the standard deviation of the mass error (*σ*_*M*_) using the median absolute deviation (MAD) method.

#### Peak identification

Recalibrated ΔMass values were binned using 0.001 Da bins to construct the ΔMass distribution. The distribution was smoothed using the median of a 7-point sliding window and then peak apexes were detected as downward zero-crossings in the first derivative of the smoothed curve. Peak widths were similarly calculated as the zero-crossing points of the second derivative; in the current version of SHIFTS they are computed only for informative purposes.

#### Peak assignation

By default SHIFTS assigns a PSM to the closest ΔMass peak if the mass deviation of the PSM from the peak falls within 3*σ*_*M*_, so that approximately 99% of PSM in each peak are assigned. This value can be user adjusted. PSM not assigned to peaks were considered as orphan PSM.

#### FDR calculation

SHIFTS calculates FDR of identification using a conventional target/decoy strategy using the corrected Xcorr score (cXcorr) (Choi and Nesvizhskii, 2008; Keller et al., 2002). A *global FDR* was calculated for each PSM as the ratio of the number of decoy PSMs to the number of target PSMs having a cXcorr equal or higher. Decoy peptides matched by Comet-PTM were observed to be almost as abundant as target peptides in the negative ΔMass region below the peak corresponding to neutral loss of Gly (Figure 2A), where ΔMass peaks were mostly produced by neutral loss of amino acids. For this reason, the global FDR was only calculated in the ΔMass region above −56 Da (Figure 2A). All the PSM are required to have FDR lower than the global FDR, without exception.

In addition, local FDR filters are also applied. Some ΔMass peaks were observed to contain an unusually high number of decoy PSM; to avoid matching false positive target PSM in these peaks, SHIFTS also calculates a *peak FDR* counting up the number of decoys and target PSM assigned to each peak, and these PSM are required to pass the peak FDR filter in addition to the global FDR filter (Figure 2C). Note that peak FDRs are often very low suggesting that the majority of PSM in these peaks are true, even when they have a low cXcorr. This happens because the probability of finding a decoy PSM in a peak by chance alone is extremely low. SHIFTS avoids matching these low scoring target PSM by applying the global FDR filter.

To apply a local filter to PSM which are not assigned to ΔMass peaks, e.g. to orphan PSM, SHIFTS models the periodic mass distribution of decoy PSM into ~1 Da-bins centered at the regions where ΔMass values concentrate, and calculates a *local FDR* by counting up decoy and target PSM in each one of these regions (Figure 2B). The local FDR filter is applied to orphan PSM in addition to the global FDR filter. Default values for peptide identification were 1% for peak and local FDR; 5% for global FDR. Among the peptides significantly changed by heteroplasmy and suspected to have relevant biological activity, including all the peptides in Table S3, only those that passed analysis with Vseq program (Cogliati et al., 2016) were considered as trustable identifications.

#### Isotopic correction

SHIFTS also performs a simple isotopic correction to minimize missassignations of the correct monoisotopic peak of the precursor. When two PSM having the same sequence are encountered having a ΔMass difference within 1 ppm of the mass difference expected for either one or two ^13^C or one ^34^S, the ΔMass of the heaviest precursor is substituted by that of the lightest one.

### Annotation of modifications

A Python in-house script was used for semisupervised annotation of the nature of peptide modifications. The script searched ΔMass values against Unimod database, taking into account the amino acid modified according to Comet-PTM output and also the preceding and consecutive residues, comparing them with the list of amino acids that could be subjected to the modification according to Unimod. If no amino acid was matched, the modification was considered as unassigned. Residues containing modifications that were fixed in the database search were considered with and without the fixed modification; when not indicated, the amino acid contains the fixed modification (e.g., C_oxidation means oxidation of carbamidomethylated Cys and K_oxidation, oxidation of TMT-labelled Lys). ΔMass values that could not be matched were tentatively tested assuming one ^13^C misassignment of the monoisotopic mass of the precursor and also as combinations of two modifications from a list of the most abundant modifications found in the corresponding proteome. Unexplained ΔMass values were termed as unknown. MS/MS fragmentation spectra from the most abundant modifications that changed their abundance in heteroplasmic mice and all the peptides in Table S3 were revised using Vseq program (Cogliati et al., 2016).

### Peptide quantification and statistical analysis

The quantitative information from TMT reporter intensities was integrated from the spectrum level to the peptide level and then to the protein level on the basis of the WSPP model (Martinez-Acedo et al., 2012; Navarro et al., 2014) using the Generic Integration Algorithm (GIA) (Garcia-Marques et al., 2016). The algorithm was modified to include into the statistical model the quantitative values of modified peptides as part of the automated workflow (Figure 3A). Briefly, for each sample *i* the values *x*_*qps*_ - log_2_ *S*_*i*_/*C* were calculated, where *S*_*i*_ is the intensity of the TMT reporter corresponding to sample *i* in the MS/MS spectrum *s* coming from peptide *p* and protein *q*, and *C* is the average intensity of all the TMT reporters from the control samples, which is used as a common reference. The log_2_-ratio of each peptide (*X*_*qp*_) was calculated as the weighted average of its spectra, the protein values (*X*_*q*_) were the weighted average of its peptides, and the grand mean 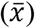 was calculated as the weighted average of all the protein values (Navarro et al., 2014). The statistical weights of spectra, peptides and proteins (*w*_*qps*_, *w*_*qp*_ and *w*_*q*_, respectively) and the variances at each one of the three levels (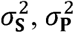, and 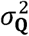, respectively), were calculated as described (Navarro et al., 2014).

The spectrum, peptide and protein variances and the protein values were firstly determined including only non-modified peptides (Figure 3B). In a second step, the modified peptides were included in the analysis, which was performed using the variances and protein values calculated previously. For each modified peptide, the standardized variable (*Z*_*pq*_) was calculated as

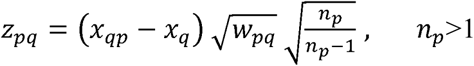

where *n*_*p*_ is the number of non-modified peptides with which the corresponding protein was quantified. *Z*_*pq*_ expresses the deviation between the peptide log_2_-ratio and the corresponding protein value in units of standard deviation. In absence of changes, the distributions of *Z*_*pq*_ followed very closely the normal distribution *N* (0,1) (Figure 3C), validating the accuracy of the model.

Significant abundance changes of modified peptides were detected by Student’s t-test comparing the *Z*_*pq*_ values of samples from heteroplasmic mice (N=4) with those of control mice (N=3) (Table S2).

### Code access

The software needed to execute the whole Comet-PTM pipeline can be downloaded from: https://github.com/CNIC-Proteomics/Comet-PTM. The software for statistical analysis of quantitative data can be downloaded from https://github.com/CNIC-Proteomics/PTM-Quant-Stats. A readme.txt file is provided with basic instructions to install and execute the package. Vseq is available upon request.

### Data Access

The data set (raw files, protein databases, search parameters and results) is available in the PeptideAtlas repository (ftp://PASS01085:TC4334fh@ftp.peptideatlas.org/), which can be downloaded via ftp.peptideatlas.org (username: PASS01085, password: TC4334fh).

## SUPPLEMENTAL FIGURE AND TABLE LEGENDS

**Figure S1.**
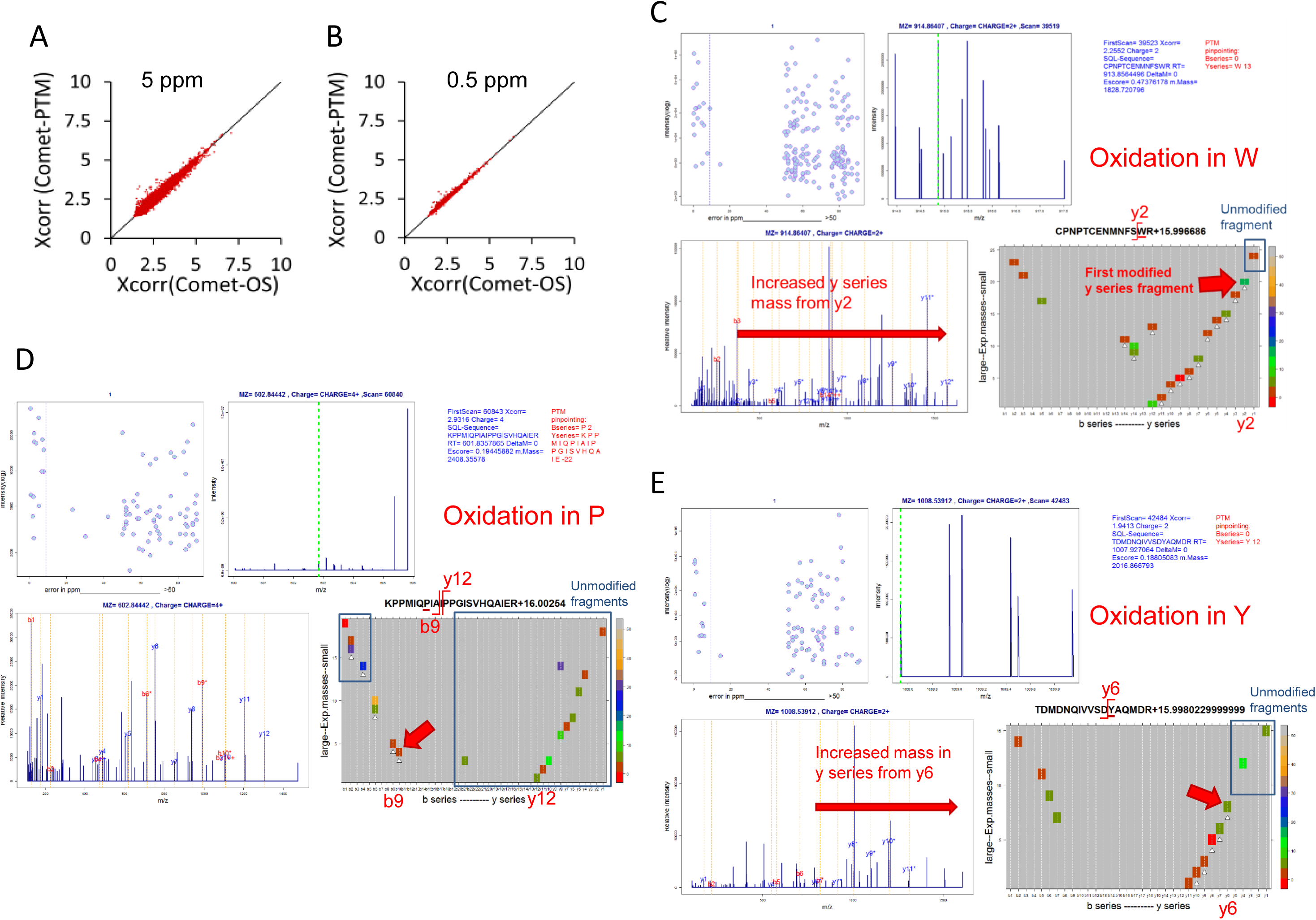
Related to Figure 1. Peptide identification performance of Comet-PTM. (**A-B**) Comparison of scores obtained from Comet-PTM and CS in the population of PSM that produced the same match with the two engines. A match was considered identical when the peptide sequence was the same and the difference between ΔMass obtained by Comet-PTM and the theoretical mass of the modification selected in CS was within 5 ppm (**A**) or 0.5 ppm (**B**). Note that the scores were practically identical, and the dispersion around the identity line was diminished when the tolerance decreased; this demonstrates that the small differences in the score are a consequence of the error in the estimation of ΔMass and not in the design of the score in Comet-PTM. (**C-E**) Vseq analyses of 3 peptides containing oxidations incorrectly assigned to Met by CS. Comet-PTM correctly assigns the oxidation to Trp (**C**) Pro (**D**) or Tyr (**E**).

**Figure S2.**
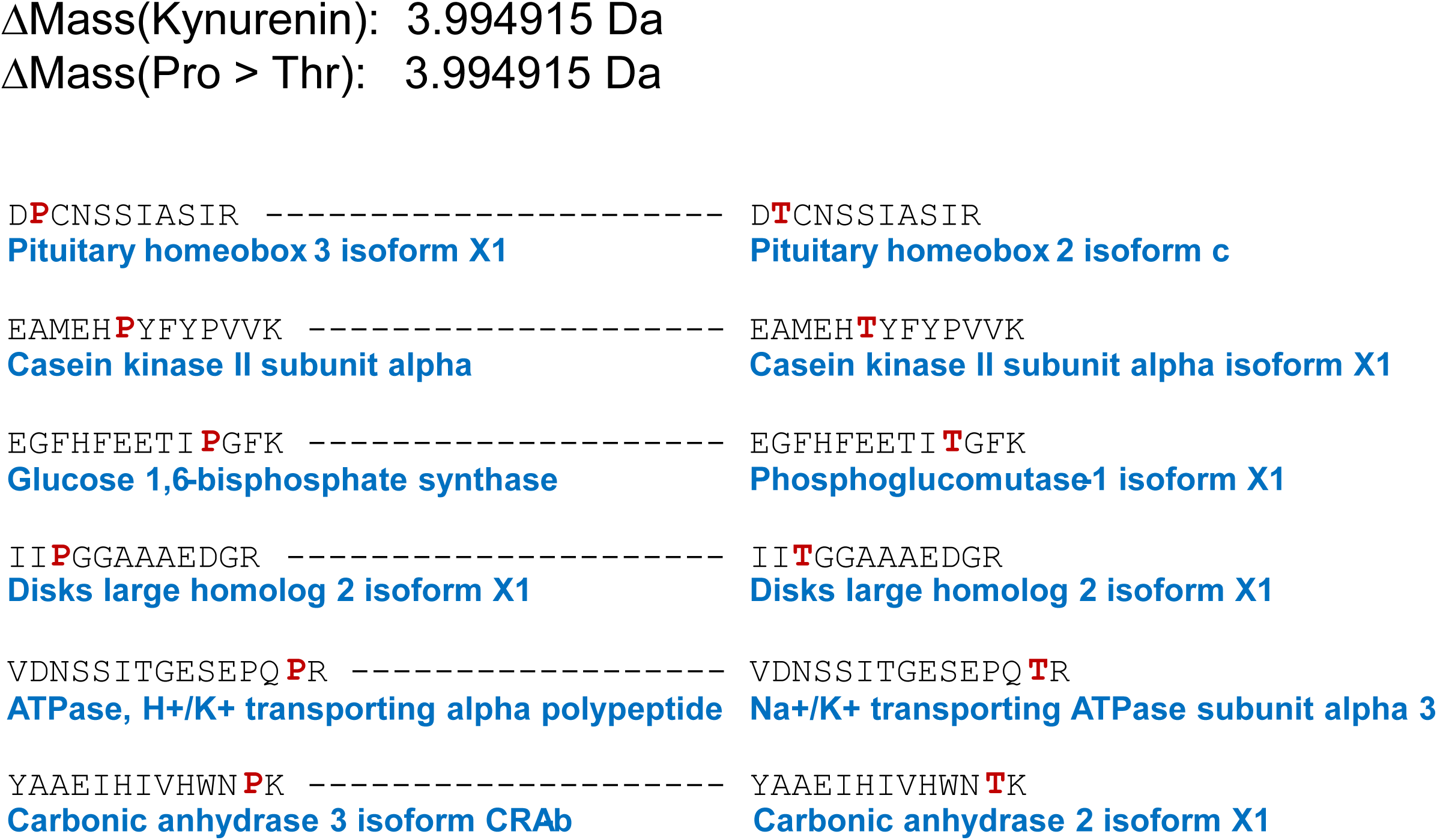
Related to Figure 2. List of peptides containing a Pro > Thr substitution assigned as a kynurenin modification in Pro. Peptide sequences together with their corresponding proteins are listed, highlighting (in red) the position of Pro and Thr residues.

**Figure S3.**
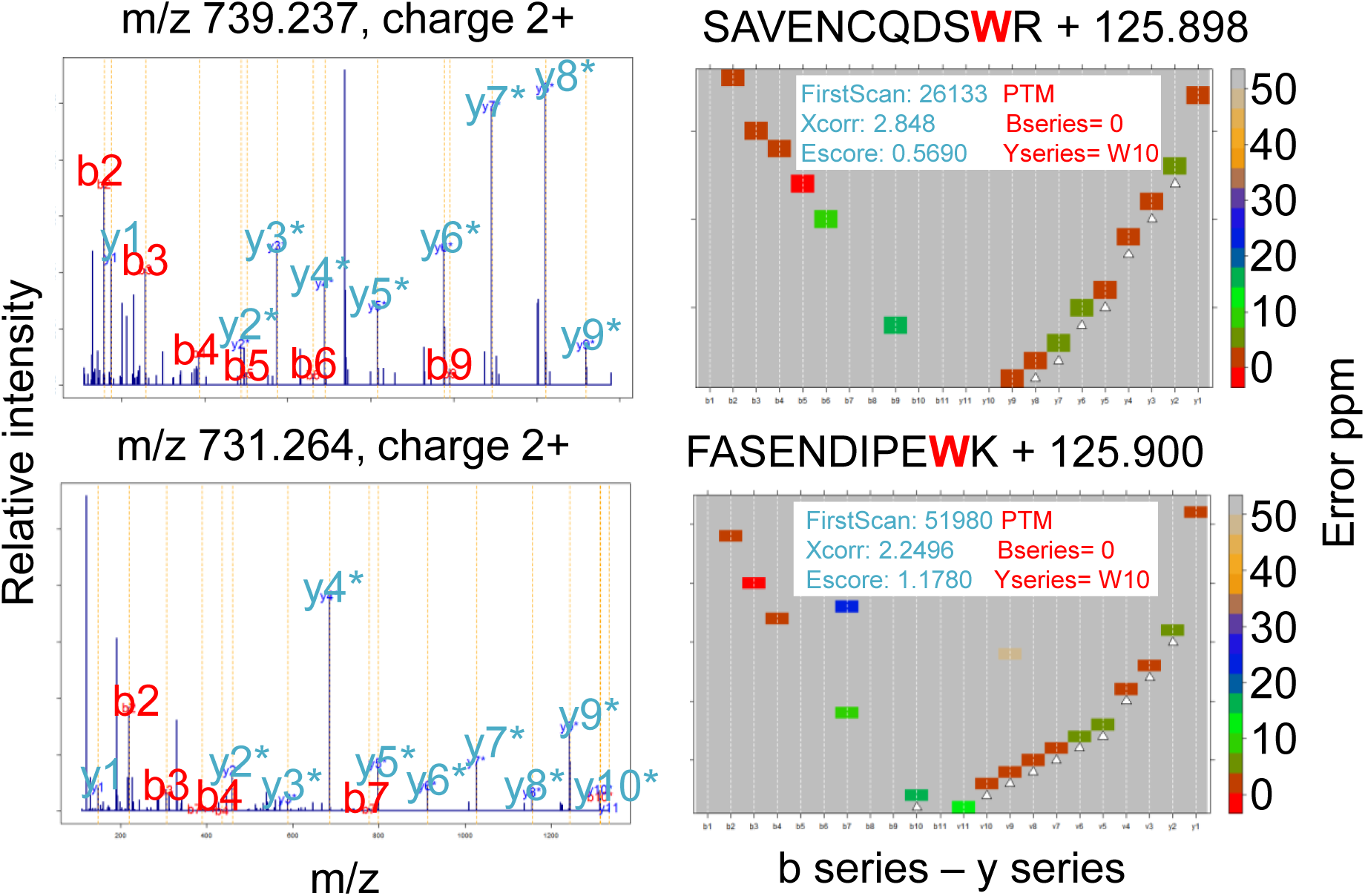
Related to Figure 2. Detection of a Trp iodination. Vseq results for 2 representative peptides containing iodinated Trp. Both the MS2 spectra (left panels) and the V-shaped heatmap distributions for the main fragmentation series (right panels), demonstrate the location of the modification in Trp.

**Figure S4.**
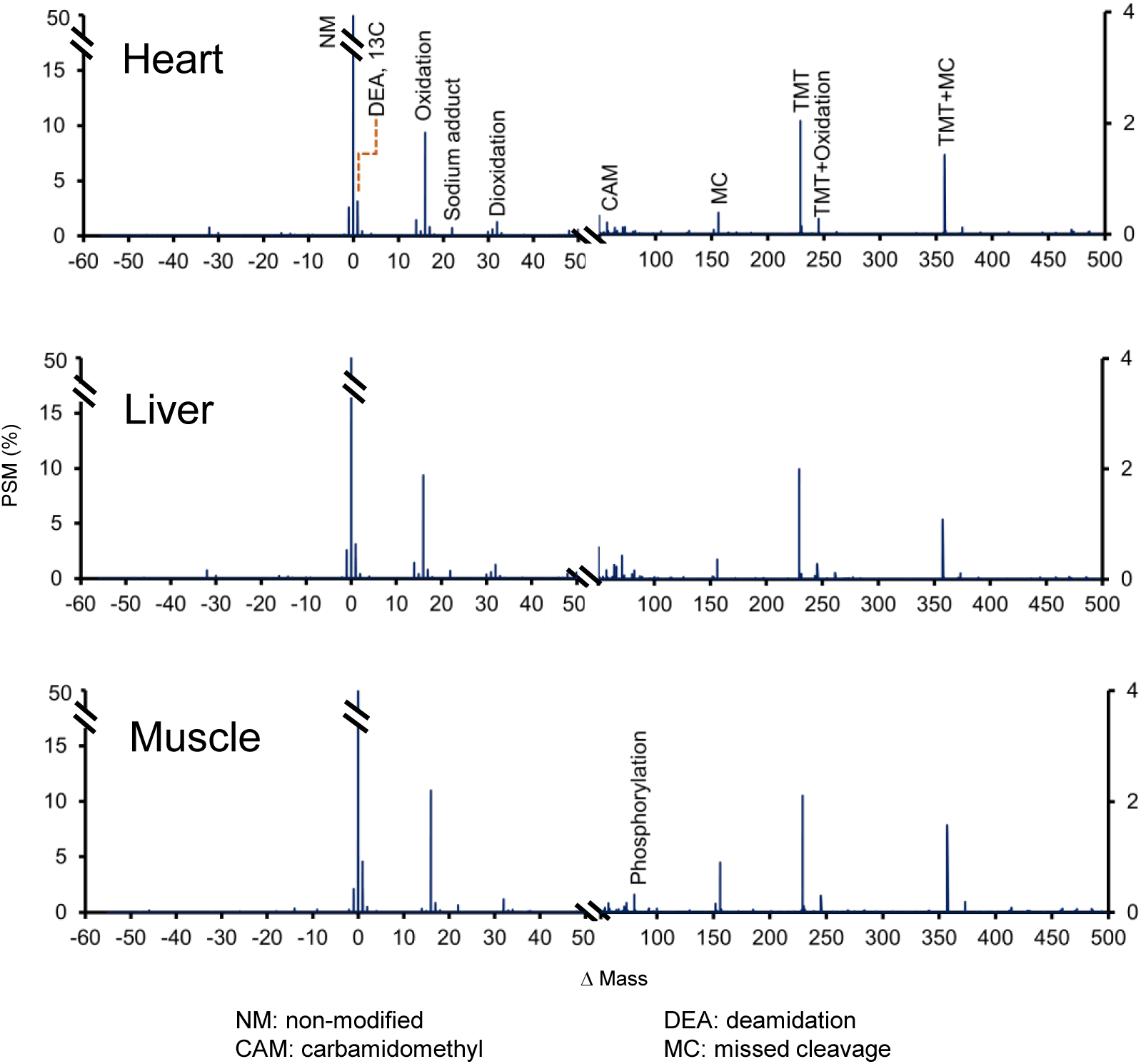
Related to Figure 4. ΔMass distribution of modified peptides identified in the 3 mouse tissues. Assignations of the most frequent ΔMass peaks are indicated.

**Figure S5.**
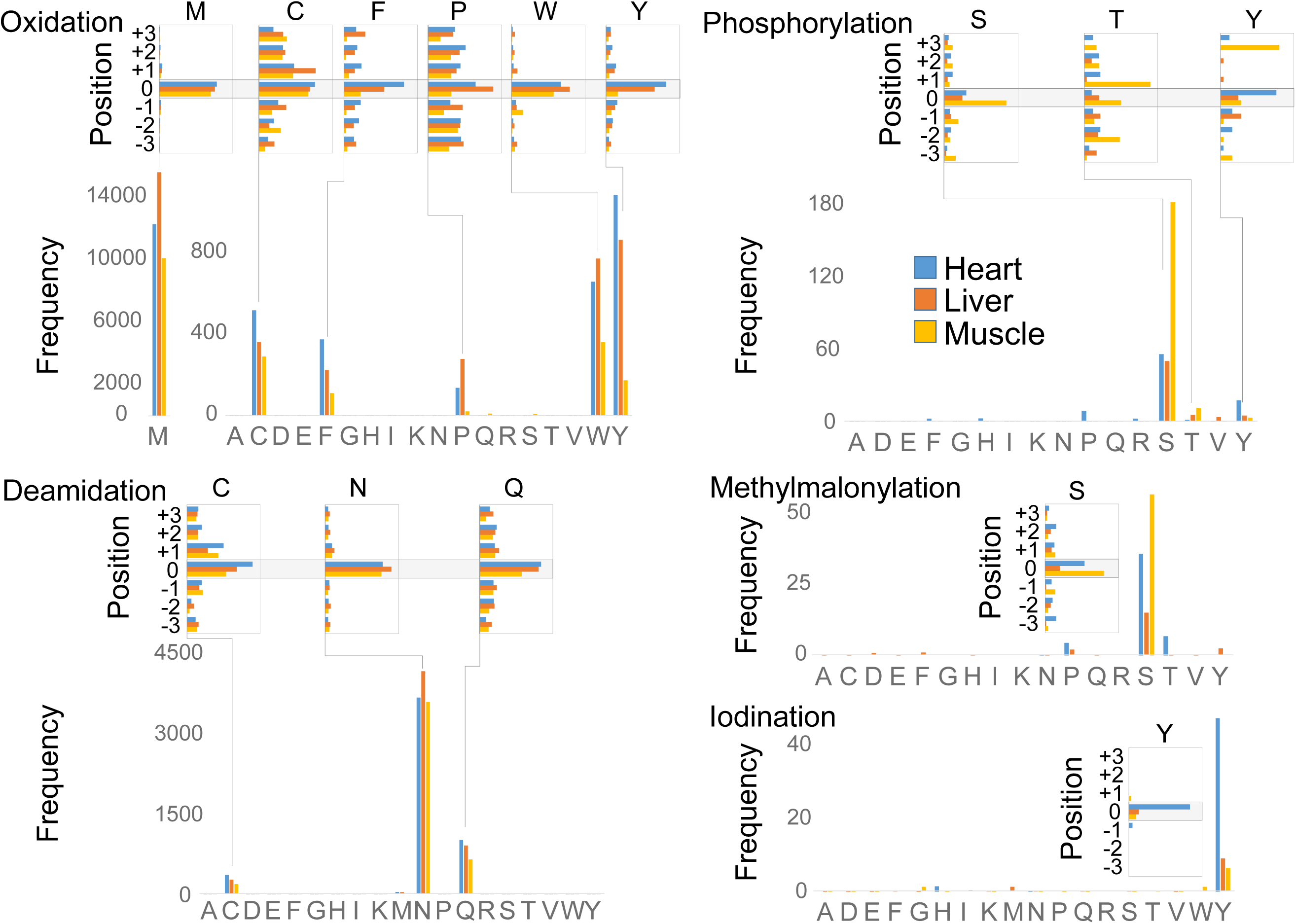
Related to Figure 4. Distribution of amino acids containing the most frequent modifications. The horizontal and vertical bar graphs have the same meaning as in Figure 2E.

**Figure S6.**
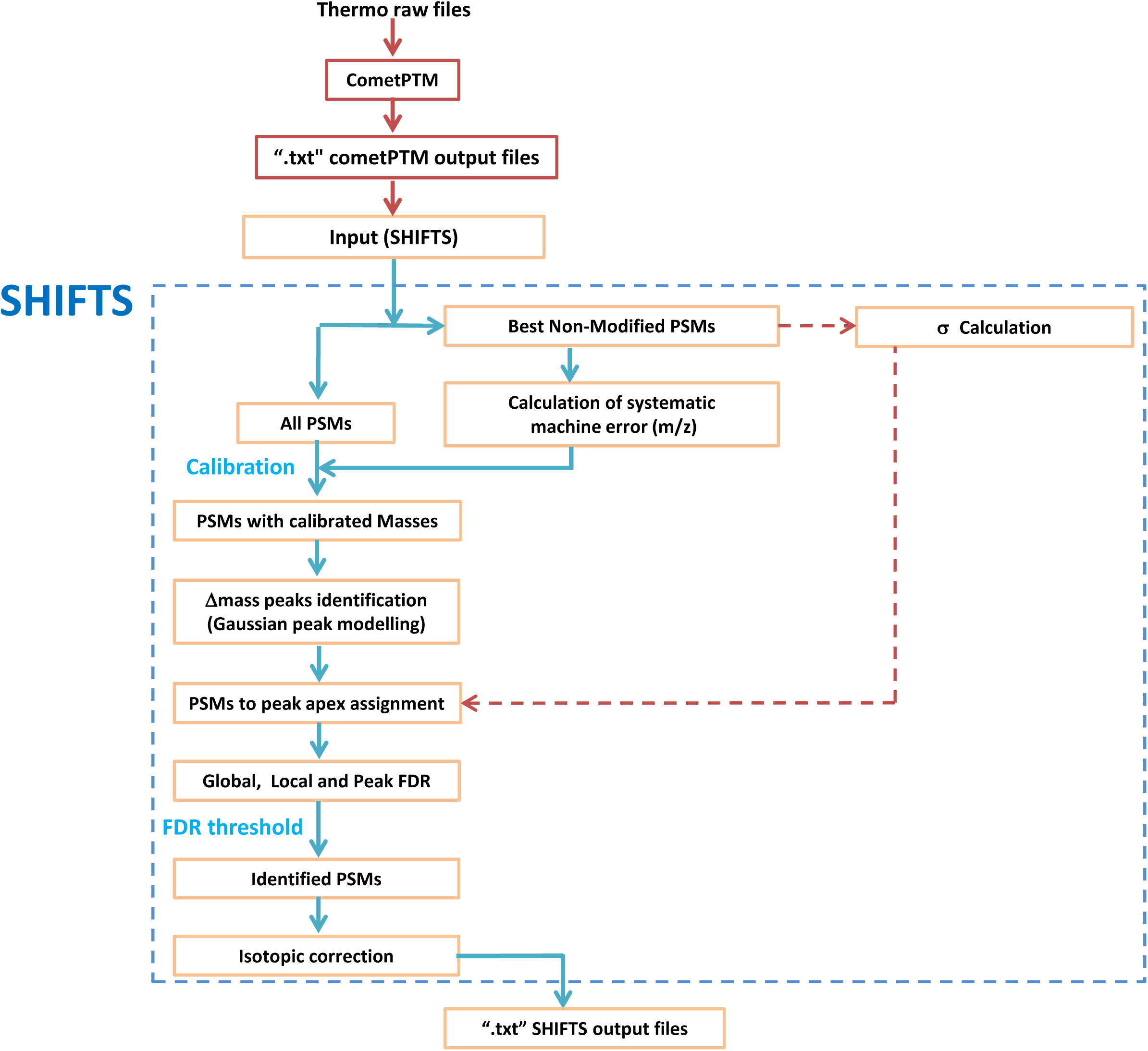
Related to Experimental Procedures. Scheme of SHIFTS algorithm. Schematic representation of SHIFTS (Systematic, Hypothesis-free Identification of modifications with controlled FDR based on ultra-Tolerant database Search), depicting how the output from Comet-PTM is processed, including mass recalibration, peak detection and FDR calculation.

**Table S1. Related to Figure 3. List of functional protein categories significantly altered by heteroplasmy (FDR < 5%) as a consequence of a coordinated protein response**

Increases in the heteroplasmic tissues, in relation to the average protein abundance of all samples are coloured in red, whereas decreases are in blue, according to the graded scale.

**Table S2. Related to Figure 4. List of modified peptides significantly altered by heteroplasmy (p < 0.05)**

Increases in peptide abundance in the heteroplasmic tissue compared to the average of peptide abundance of all samples are coloured in red, whereas decreases are in blue, according to the graded scale. p-values are calculated using Student’s t-test.

**Table S3. Related to Figure 6. Conservation and structural analysis of heteroplasmy-modified peptides of OXPHOS complexes I, III, IV and V.**

The residues harboring the modification are underlined. Residues predicted to be functional (conservation score 8-9 and exposed) are highlighted in red; residues predicted to have structural implications (conservation score 9 and buried), in blue; highly conserved residues (conservation score >5), in green. Most PTMs are on highly conserved residues or next to highly conserved residues. Conservation scores were rated from 9 (highest conservation) to 1 (highest variability) by the ConSurf server (Ashkenazy et al., 2016).

